# NMNAT2-SARM1 Axis Drives Redox Failure and Disrupts APP Processing in Neurons

**DOI:** 10.64898/2026.04.16.718990

**Authors:** Andrea Enriquez, Sen Yang, Karen Ling, Paymaan Jafar-Nejad, Hui-Chen Lu

## Abstract

Metabolic dysfunction and proteinopathy are hallmarks of many neurodegenerative diseases, yet their mechanistic interplay remains poorly understood. Here, we demonstrate that amyloid precursor protein (APP) processing in cortical neurons is disrupted upon loss of Nicotinamide mononucleotide adenylyltransferase 2 (NMNAT2), the NAD⁺-synthesizing enzyme in neurons, resulting in accumulation of APP C-terminal fragments (APP-CTFs). Knockdown (KD) of the NAD⁺ hydrolase sterile alpha and TIR motif-containing protein 1 (SARM1) restores APP-CTF levels in NMNAT2 knockout (KO) neurons to wild-type levels, whereas NAD⁺ supplementation yields modest rescue. Redox profiling indicates that NMNAT2 loss reduces NAD⁺/NADH redox potential when APP-CTF starts accumulating. Seahorse metabolic flux analysis shows that NMNAT2 deficiency induces early glycolytic impairment, followed by deficits in mitochondrial respiration. Notably, SARM1 KD, but not NAD⁺ supplementation, rescues mitochondrial function in NMNAT2 KO neurons. Temporal profiling of NMNAT2 KO neurons revealed a biphasic pattern in APP-CTF accumulation, with an initial gradual increase followed by a marked acceleration, paralleling the transition from an initially small number to a substantially greater number of differentially expressed proteins. Pathway enrichment analysis of proteomic changes suggests JNK/MAPK signaling is upregulated in the early phase, with late-phase downregulation of mitochondrial function and upregulation of endoplasmic reticulum stress and unfolded protein response pathways. Collectively, these findings demonstrate that neuronal NAD⁺ depletion drives a progressive, SARM1-dependent disruption of redox homeostasis and proteostasis, resulting in impaired APP processing. The NMNAT2–SARM1 axis emerges as a critical pathway linking metabolic stress to proteinopathy, positioning SARM1 as a key mediator of neurodegenerative dysfunction.

## Introduction

Neurodegenerative diseases involve system-level dysfunctions that are caused by a variety of environmental and genetic factors. FDG-PET imaging studies have revealed progressive cerebral glucose hypometabolism in patients with Alzheimer’s disease (AD), Parkinson’s disease (PD), and Huntington’s disease (HD) (1–4). Their pathological features include progressive, age-dependent proteinopathy and neuronal loss (5–7), and growing evidence suggests that aging-associated declines in cellular energy metabolism contribute to the onset and progression of neurodegeneration (8–11). Amyloid β (Aβ) deposition examinations in AD patients have shown that brain regions with Aβ plaques are also areas with significant hypometabolism of glucose (12). Currently, the mechanistic link between glucose hypometabolism and proteinopathy remains elusive.

Nicotinamide adenine dinucleotide (NAD⁺) occupies a central position at the interface of energy metabolism (13), redox balance (14), and pathways involved in signaling cascades that regulate proteostasis (15). Protein processing, which includes protein folding, post-translational modifications, trafficking through organelles, and proteasomal clearance, involves a wide array of molecular mechanisms and organelles and is particularly sensitive to metabolic stress (16–19). Nicotinamide mononucleotide adenylyl transferase 2 (NMNAT2) is the most abundant NAD^+^-synthesizing enzyme in mammalian cortical neurons (20–22). Its expression is decreased across various proteinopathies, such as AD, PD, HD (23), and in the spinal cord of amyotrophic lateral sclerosis patients (24). Our recent studies show that deleting NMNAT2 from glutamatergic neurons in mice results in defective glucose metabolism and several neurodegenerative phenotypes, including axonal amyloid precursor protein (APP) accumulations and neurodegeneration in a sterile alpha and TIR motif-containing protein 1 (SARM1)-dependent manner (25, 26). SARM1 is an NAD^+^ hydrolase that is activated when NMNAT2 levels are reduced (27), leading to NAD⁺ depletion and axonal degeneration (28).

APP is a transmembrane protein that is primarily cleaved by two canonical pathways: non-amyloidogenic APP processing or amyloidogenic APP processing, both of which are driven by α-, β-, and γ-secretases (29). The cleavage by α- and β-secretases generates APP C-terminal fragments (APP-CTFs), and subsequent cleavage by γ-secretases leads to the production of Aβ. Significant work has focused on abnormal APP cleavage for neurotoxic Aβ generation (30). However, there is growing evidence highlighting the toxic effects of APP-CTFs, including decreased mitochondrial respiration, impaired endolysosomal dysfunction, synapse loss, and neuronal hyperexcitability mitophagy (31–36). Bretou et al. (37) showed that both α- and β-derived CTFs reproduce endolysosomal dysfunction, indicating that toxicity derives from APP-CTF signaling rather than from a specific cleavage product. Additionally, intracellular accumulation of β-CTFs in human neurons directly impairs lysosomal function (38). Studies with AD mouse models show that β-CTF accumulation contributes directly to synaptic dysfunction and non-cognitive behavioral alterations (36). Interestingly, CTFs could be released with exosomes upon neuronal lysosomal dysfunction (39). The proteolytic processing of APP can be altered by disrupted intracellular trafficking, since secretases are localized in different cellular compartments, and changes in APP trafficking can affect its access to these enzymes (40–42). Moreover, defective energy homeostasis has also been shown to alter APP processing (43–45).

Our previous findings of axonal APP accumulation in NMNAT2 knockout (KO) neurons were revealed using an APP antibody that recognizes the C-terminal epitope. Thus, it is plausible that APP-CTFs are increased in NMNAT2 KO axons (25). Here, we investigated whether NMNAT2 loss alters APP processing and results in APP-CTF accumulation. A combination of biochemical, metabolic assays, pharmacological/antisense oligo manipulations, and proteomic profiling was conducted. Our data suggest that NMNAT2 loss induces mechanistically separable yet interconnected proteostatic and metabolic defects that lead to a progressive increase in APP-CTFs, and that the NMNAT2-SARM1 axis in cortical neurons plays an important role in regulating APP proteomic processing.

## Methods

### Mice

Blad mice with a NMNAT2 null mutation were generated by transposon-mediated gene-trap mutagenesis as previously described (46). Both male and female mice were used throughout this study. All mice were housed under standard conditions with ad libitum food and water and maintained on a 12-hour light/dark cycle. Animal care and experimental procedures were carried out in compliance with the NIH Guidelines for the Care and Use of Laboratory Animals and approved by the Institutional Animal Care and Use Committees at Indiana University.

### Genotyping

Genotyping lysates were prepared by placing embryonic tail samples in digestion buffer (50 mM KCl, 10 mM Tris–HCl, 0.1% Triton X-100, 0.2 mg/ml proteinase K, pH 9.0) and incubating at 60 °C for 15 minutes in a thermal shaker set to 1500rpm. Samples were then incubated at 95 °C for 10 minutes to denature the proteinase K and centrifuged at max speed in a microcentrifuge for 5 minutes. The supernatants were used as DNA templates for polymerase chain reactions using the EconoTaq® Plus Green 2X (Biosearch™ Technologies) master mix or the QIAGEN® Fast Cycling PCR Kit. Samples were loaded onto a 1.6% agarose gel and ran at 135 V for 30 minutes. The primer sequences used for genotyping have been previously described (25).

### Primary cortical neuron culture

Primary cortical mouse neurons were prepared using E14.5–16.5 embryos from heterozygous (het) NMNAT2-Blad mouse matings. Cortices of both male and female embryos were dissected and genotyped, to then be sorted into wild-type (WT) and KO groups. Following dissection and genotyping, the pooled tissue was prepared for plating using the Papain Dissociation System (Worthington®) following the manufacturer’s protocol. Cells were counted using a hemocytometer and plated at a density of 1.43×10^5^ cells/cm^2^. Primary neurons were maintained in Neurobasal Media (Gibco™) supplemented with B-27 (Gibco™), GlutaMAX™ (Gibco™), and penicillin–streptomycin (Gibco™). All cell culture preparations were performed under sterile conditions, and cells were incubated at 37 °C with 5% CO^2^ and appropriate humidity. One-third of the media was replenished every 3 days. Samples were collected at day-in-vitro (DIV) indicated in Results.

### Western Blotting

Neurons were lysed using 150ul of RIPA buffer per well [20 mM of Tris-base, 150 mM of NaCl, 2 mM EDTA, 1% Triton X-100, 0.5% sodium deoxycholate, 0.1% sodium dodecyl sulfate (SDS), 1x cOmplete™ Mini protease inhibitor cocktail (Roche®) and phosphatase inhibitor cocktail 3 (Sigma-Aldrich® P0044), pH 7.4] and homogenized for 15 seconds with a pestle motor using disposable pellet pestles. Samples were then sonicated using a VMR Branson Sonifier 250 for 10 intervals set at duty cycle 30/output 3 and stored at -80°C.

Protein concentration was measured and normalized using the Pierce™ BCA Protein Assay Kit (Thermo Scientific™) and samples were prepared in Laemmli SDS buffer. 20ul of protein samples were loaded onto a 16.5% SDS–polyacrylamide gel and run at 80 volts (V) for 20 min and 110V until the lower band of loading dye reached the bottom of the gel. Protein lanes were transferred to a nitrocellulose membrane with a 0.22 μm pore size using a tank transfer system (Bio-Rad). For sample normalization, nitrocellulose membranes were stained using Ponceau S for 30 seconds, rinsed 3x with Milli-Q® dH2O, and imaged using a Gel Doc™ XR+ imager. Membranes were blocked for 1 hour at room temperature with blocking solution (1:1 BlockOut® Universal Blocking Buffer and 0.1% TBST), followed by a 24-hour incubation with APP primary antibody (ab32136, abcam) diluted in blocking solution. The next day, membranes were washed with 0.1% TBST (3 x 10min) followed by a 2-hour incubation with the secondary antibody IRDye 680 LT (92-68021, Li-COR) diluted in blocking solution. Membranes were then washed with 0.1% TBST (3 x 10min) and stored in TBS until imaging. Western blot images were acquired with a LI-COR Odyssey® scanner and software (LICORbio™), and protein abundance was quantified using the NIH Fiji ImageJ software (47). Two quantification methods were used to analyze blots in this study. Total expression band values were obtained using the Fiji Gels built-in analysis function. Briefly, same size boxes were placed on top of each band and lane intensity was plotted using the Fiji Plot Lanes function. Area under the curve was then extracted from each plot, and intensity was normalized to WT controls in each gel. For the intensity signal curve plots, all gel sizes were matched by resizing the distance between the 25 and 10 kilodalton (kDa) molecular weight markers to 200 pixels using the Resize function. Once gels were resized, a line of equal length was placed at the 20kDa protein marker and positioned at the center of the bands signal for each sample. The Fiji Plot Profile function was used to create response curves of signal intensity at each point of the line, and signal across this set distance was acquired for each sample. Signal intensity was normalized to the WT controls in each gel.

### BACE1 Inhibitor Treatment

β-Secretase Inhibitor IV (Cayman Chemical, 23388), a known inhibitor for β-site amyloid protein cleaving enzyme 1 (BACE1), was dissolved in DMSO to a stock concentration of 1mM and stored at -20° C. For experiments, the inhibitor was diluted in cell culture medium to a working concentration of 3 µM. At DIV10, cells were treated with either DMSO or a BACE1 inhibitor (B-Inh) by replacing one-third of the medium in each well, resulting in a final concentration of 1 µM. Cells were collected at DIV13 for Western blotting.

### NAD^+^ supplementation

NAD^+^ (Roche, NAD100-RO) was dissolved at a stock concentration of 100mM in 1x PBS, filtered with a 0.22 µM syringe filter, and stored at -80 °C for up to 7 days. On the first day of supplementation, NAD^+^ was added to the culture medium at a final concentration of 1mM. The same volume of PBS was added to the Neurobasal medium to use as a control. In the following days, one-third of the culture medium was refreshed daily with 1 mM NAD^+^ or PBS until cells were collected.

### Antisense Oligonucleotide Treatment

Antisense oligonucleotides (ASOs) used in this study have been previously described and validated (25). A non-targeting ASO, 5’-CCTATAGGACTATCCAGGAA-3’ (Ctrl ASO) was used as a control. An ASO targeting mouse SARM1 mRNA, 5’-GGTAAGAGCCTTAGGCACGC-3’ (SARM1 ASO) was used to knockdown (KD) SARM1 expression. To start treatment, ASOs were diluted in neuronal culture media and added to neurons at a final concentration of 5 µM, and one-third of the culture medium was refreshed every 3 days with 5uM concentration of ASOs.

### NAD^+^/NADH-Glo™ bioluminescent assay

NAD^+^/NADH measurements were carried out as previously described (25). Briefly, WT and KO neurons were cultured in 96-well plates at a density of 1.3×10^3^ cells/mm^2^. Following Promega’s NAD+/NADH-Glo™ Assay protocol, DIV8 or DIV12 neurons were lysed in DTAB base buffer (100 mM Sodium Carbonate, 20 mM Sodium Bicarbonate, 10 mM Nicotinamide, 0.05% TritonX-100, pH 10–11). Lysates were then split for total NAD⁺+NADH and NADH-only measurements, with selective degradation of NAD⁺ in the NADH samples prior to detection. The Promega detection reagent was then added to all wells, and an enzymatic cycling reaction generates a luminescent signal proportional to NADH content. Luminescence was measured using a microplate reader, and NAD⁺ levels were calculated by subtracting NADH from total values. NAD^+^ and NADH measurements were normalized to protein concentration measured with Pierce™ BCA Protein Assay Kit (Thermo Scientific™).

### Seahorse XF Glycolytic Rate Assay

Cellular glycolysis was measured using the Seahorse XF Glycolytic Rate Assay Kit (Agilent, 103344–100) following the manufacturer’s protocol. Primary cortical neurons were prepared as described above. Neurons were seeded in Seahorse XF24 Cell Culture Microplates at a density of 40,000 cells per well in 250ul of XF DMEM medium (Agilent, 103575-100). Assays were performed on an XFe24 Seahorse Analyzer (Agilent Technologies) at DIV8 and DIV12. Following basal measurements, the medium is injected with 15 µM N-methyl-D-aspartic acid (NMDA) and 2 µM Glycine. Cells are sequentially treated with the mitochondrial inhibitor rotenone/antimycin A (Rot/AA) at 0.5µM, and glycolysis inhibitor 2-deoxyglucose (2DG) at 50mM. Extracellular acidification rate (ECAR) and oxygen consumption rate (OCR) are recorded and used to calculate glycolytic proton efflux rate (glycoPER). GlycoPER was analyzed using Wave software (Agilent Technologies) following the manufacturer’s instructions. Data were normalized to total protein content per well using the Pierce™ BCA Protein Assay Kit (Thermo Scientific™).

### Seahorse XF Cell MitoStress Test

Mitochondrial function was measured using the Seahorse XF Cell MitoStress Test Kit (Agilent, 103015–100) following the manufacturer’s protocol. Primary cortical neurons were prepared as described above. Neurons were seeded in Seahorse XF24 Cell Culture Microplates at a density of 40,000 cells per well in 250ul of XF DMEM medium (Agilent, 103575-100). Assays were performed at DIV8 and DIV12 using an XFe24 Seahorse Analyzer (Agilent Technologies). Sequential injections of mitochondrial modulators were performed at the following final concentrations: 1.5 µM oligomycin, 2 µM carbonyl cyanide-4 (trifluoromethoxy) phenylhydrazone (FCCP), and 0.5 µM Rot/AA. OCR was recorded and analyzed using Wave software (Agilent Technologies) following the manufacturer’s instructions. Data were normalized to total protein content per well using the Pierce™ BCA Protein Assay Kit (Thermo Scientific™).

### Proteomic profiling

WT control and KO neurons were plated on 12-well plates as described above. At DIV8 or 12, cells were rinsed 3x with ice-cold PBS. 300ul of PBS was added to each well and cells were detached from the plate using a sterile cell scraper. 2 wells were combined into 1 sample and centrifuged at 300g for 8 minutes. Cell pellets were rinsed once with ice-cold PBS and centrifuged again at 300g for 6 minutes. Samples were flash frozen in liquid nitrogen and stored at -80° C.

Liquid chromatography–mass spectrometry (LC–MS) experiments, along with preliminary data normalization and analysis, were conducted at the Center for Proteome Analysis at Indiana University School of Medicine (Indianapolis, IN, USA). Gene set enrichment analysis was performed using Enrichr (https://maayanlab.cloud/Enrichr/) and STRING (https://string-db.org). Proteins with significant p-values were separated into upregulated and downregulated groups, as well as by treatment conditions. Protein lists were imported into Enrichr and Gene Ontology (GO) Biological Processes (BP) were extracted. Data lists were also uploaded onto STRING to create full confidence string networks using databases, experiments, and co-expression as active interaction sources, with a minimum required interaction score of 0.4. Clustering was done by using the DBSCAN with an epsilon parameter of 4, and top GO terms for BP, molecular function (MF), and cellular compartment (CC) terms were extracted by enrichment analysis.

### Statistical Analyses

All data sets were tested for normality of residuals using the D’Agostino-Pearson normality test. For data sets with normally distributed residuals, we used one-way ANOVA with Tukey’s multiple comparison test, two-way ANOVA with Tukey’s multiple comparison test, or unpaired student’s t-test. For data sets that did not pass the D’Agostino-Pearson normality test, one-way ANOVA was replaced by Kruskal–Wallis test & Dunn’s multiple comparison test; two-way ANOVA was replaced by Multiple Mann–Whitney test or re-formatted for Kruskal–Wallis test; unpaired student’s t-test was replaced by the Mann–Whitney test. The criterion for statistical significance was set at p < 0.05 for all statistical analyses.

## Results

### Neuronal NMNAT2 loss results in SARM1-dependent APP-CTF accumulations

The cleavage of full-length APP (APP-FL) by an α-secretase yields a secreted-APP α (sAPP-α) fragment and a C-terminal fragment 83 (CTF-83) (29) (Figure 1A). The subsequent cleavage of CTF-83 by γ-secretase results in P3 and the APP Intracellular Domain (AICD). When APP-FL gets cleaved by β-secretase, it produces a secreted-APP β (sAPP-β) fragment and a C-terminal fragment 99 (CTF-99). The CTF-99 is then cleaved by γ-secretase to yield Aβ peptide and an AICD. To evaluate the effects of NMNAT2 loss on APP processing, we first performed Western blotting analyses to quantify APP-FL and APP-CTFs in DIV12 primary control and NMNAT2 KO neurons using a C-terminal targeting APP antibody (Figure 1B). We found that NMNAT2 KO neurons exhibit 2-fold higher expression of APP-CTFs, ranging from 10-18 kDa, and ∼20% less APP-FL than WT neurons (Figure 1C). The cleaving sites of APP processing secretases only differ by a few amino acids (29), making it challenging to clearly separate fragment products by size in Western blots. Thus, we conducted LC-MS proteomics on DIV12 WT and NMNAT2 KO samples to determine the abundance of CTF-83 and CTF-99 by searching peptide sequences that match the α- and β-secretase cleavage sites. These analyses found that both CTF-83 and CTF-99 corresponding peptides were significantly increased in DIV12 NMNAT2 KO neurons (Figure 1E).

**Figure 1.**
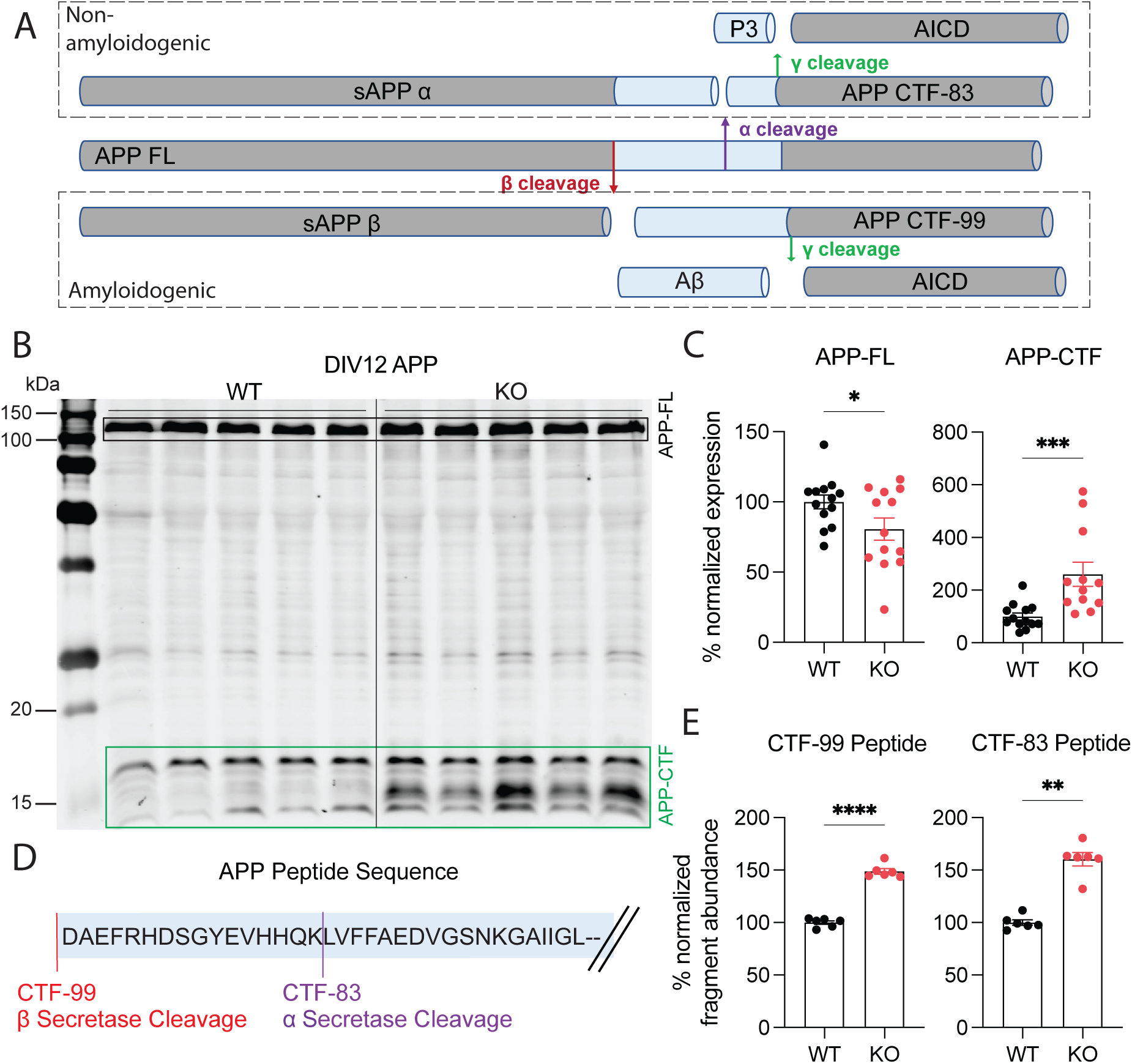
(A) Schematic diagram of non-amyloidogenic and amyloidogenic APP processing pathways. (B) Western blot of lysates from DIV12 wild-type (WT) and NMNAT2 knockout (KO) primary mouse cortical neuron cultures show APP full-length (APP-FL) and APP C-terminal fragments (APP-CTF). (C) Western blot quantification analysis. Intensity of APP-FL band and APP-CTF cluster of bands relative to WT neurons. DIV12; APP-FL n=13 WT, 13 KO; APP-CTF n=13 WT, 12 KO. APP-FL data analyzed with unpaired t-test and APP-CTF data analyzed with Mann-Whitney test. *, p=0.049, ***, p=0.0002. Samples were collected from 3 independent experiments. (D) Peptide sequence of APP illustrating the location at which beta and alpha secretases cleave the protein. (E) LC-MS/MS quantification of peptide sequences for the CTF-99 fragment as a result of beta-secretase cleavage and CTF-83 fragment as a result of alpha-secretase cleavage. DIV12; n=6 WT, 6 KO; CTF-99 data was analyzed with unpaired t-test and CTF-83 data analyzed with Mann-Whitney test. **, p=0.0022, ****, p<0.0001. Samples were collected from 2 independent experiments. All bar graphs represent mean ± SEM. Figure 2

To further confirm the increase of CTF-83 and CTF-99, we treated WT and NMNAT2 KO neurons with 1µM β-Secretase Inhibitor IV, a well-characterized BACE1 inhibitor (48, 49), from DIV10-13 to examine whether the band corresponding to CTF-99 is decreased in DIV13 NMNAT2 KO neurons (Supp. Figure 1A). We found that BACE1 inhibition increases APP-FL in both WT and KO-treated samples (Supp. Figure 1B), suggesting that BACE1 is active in DIV10-13 cortical neurons and that the inhibitor is effective. BACE1 inhibition also significantly reduced CTF-99 abundance (Supp. Figure 1D) while increasing CTF-83 levels in KO neurons (Supp. Figure 1E). However, BACE1 inhibition did not reduce the abundance of 10-18kDa APP-CTFs in KO neurons compared to the vehicle-KO group (Supp. Figure 1C). These observations allow us not only to verify the locations of CTF-83 vs CTF-99 on Western blots, but also to rule out BACE1’s contribution to the CTF increases observed in NMNAT2 KO neurons.

APP accumulation phenotypes in NMNAT2 KO neurons depend on SARM1 (25, 26). Loss of SARM1 has been reported to decrease amyloidogenic protein aggregation, supporting a connection between SARM1-driven metabolic stress responses and protein homeostasis (50). To determine whether SARM1 KD reduces APP-CTF accumulations in NMNAT2 KO neurons, we treated WT and KO neurons with Ctrl- or SARM1-ASO from DIV1-12 as previously described (25) (Figure 2A). We found that APP-CTF levels in SARM1-ASO-treated KO neurons were restored to WT levels while APP-FL levels remained unchanged (Figure 2B-C). Surprisingly, 1mM NAD^+^ supplementation during DIV5-12 only partially reduces APP-CTF levels in NMNAT2 KO neurons (Figure 2D-F).

**Figure 2.**
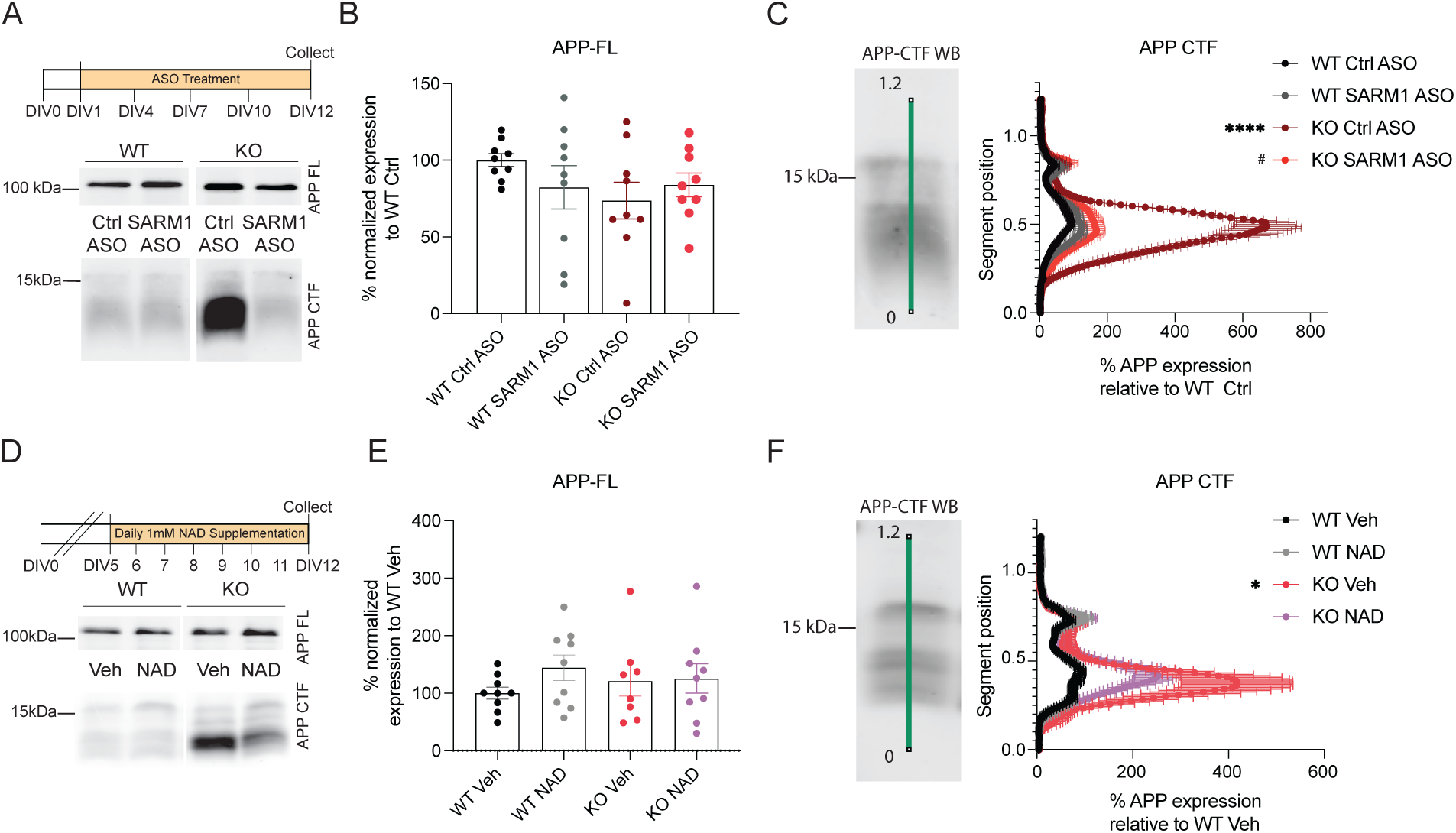
(A) Western blot of APP-FL and CTF in WT and KO neurons treated with Ctrl ASO or SARM1 targeting ASO. (B) Quantification of APP-FL band expression relative to WT Ctrl ASO. Data was analyzed using one-way ANOVA with Tukey’s multiple comparisons test. (C) Quantification of band intensity across APP-CTFs. Data was analyzed using Kruskal-Wallis test with Dunn’s multiple comparisons test. WT Ctrl ASO vs. KO Ctrl ASO ****, p=<0.0001. KO Ctrl ASO vs. KO SARM1 ASO ^#^, p=0.0197. ****, p<0.0001. n=9 WT Ctrl ASO, 9 WT SARM1 ASO, 8 KO Ctrl ASO, 9 KO SARM1 ASO. (D) Western blot of APP-FL and CTFs in control and NAD treated WT and KO neurons. (E) Quantification of APP-FL band expression relative to WT control. Data was analyzed using one-way ANOVA with Tukey’s multiple comparisons test. (F) Quantification of APP-CTFs band intensity relative to WT control. Data was analyzed using Kruskal-Wallis test with Dunn’s multiple comparisons test. WT Veh vs. KO Veh *, p=0.0132. n=9 WT, 9 WT+NAD, 8KO, 9 KO+NAD. Samples were collected from 3 independent experiments. Samples were collected from 3 independent experiments. All bar graphs represent mean ± SEM.

### SARM1 KD does not restore NAD^+^ Levels in NMNAT2 KO neurons to WT levels

SARM1, a metabolic sensor activated by a reduced NAD⁺/NMN ratio following NMNAT2 loss, promotes further NAD⁺ depletion and axonal degeneration (27, 28). Our prior studies identified DIV8 as the earliest stage when axonal deficits emerge in NMNAT2 KO neurons (25). We therefore quantified NAD^+^/NADH ratios at DIV8 and DIV12 NMNAT2 KO and WT neurons (Figure 3A). Consistent with prior observations, NMNAT2 KO neurons treated with control ASO exhibited significantly reduced NAD⁺, NADH, and NAD⁺/NADH ratios compared to WT at DIV8 (Fig. 3B–D). SARM1 ASO treatment partially corrected this redox imbalance, restoring NADH levels and the NAD⁺/NADH ratio to near-control levels, but without fully rescuing NAD⁺ abundance (Fig. 3B–D). At DIV12, despite significant increases in NAD^+^ and NADH levels in NMNAT2 KO neurons with SARM1 ASO than KO neurons with control ASO, NAD^+^ was not restored to control levels (Fig. 3E-G). Together, these data indicate that SARM1 knockdown mitigates NAD⁺ depletion in NMNAT2-deficient neurons but is insufficient to fully restore NAD⁺ homeostasis.

**Figure 3.**
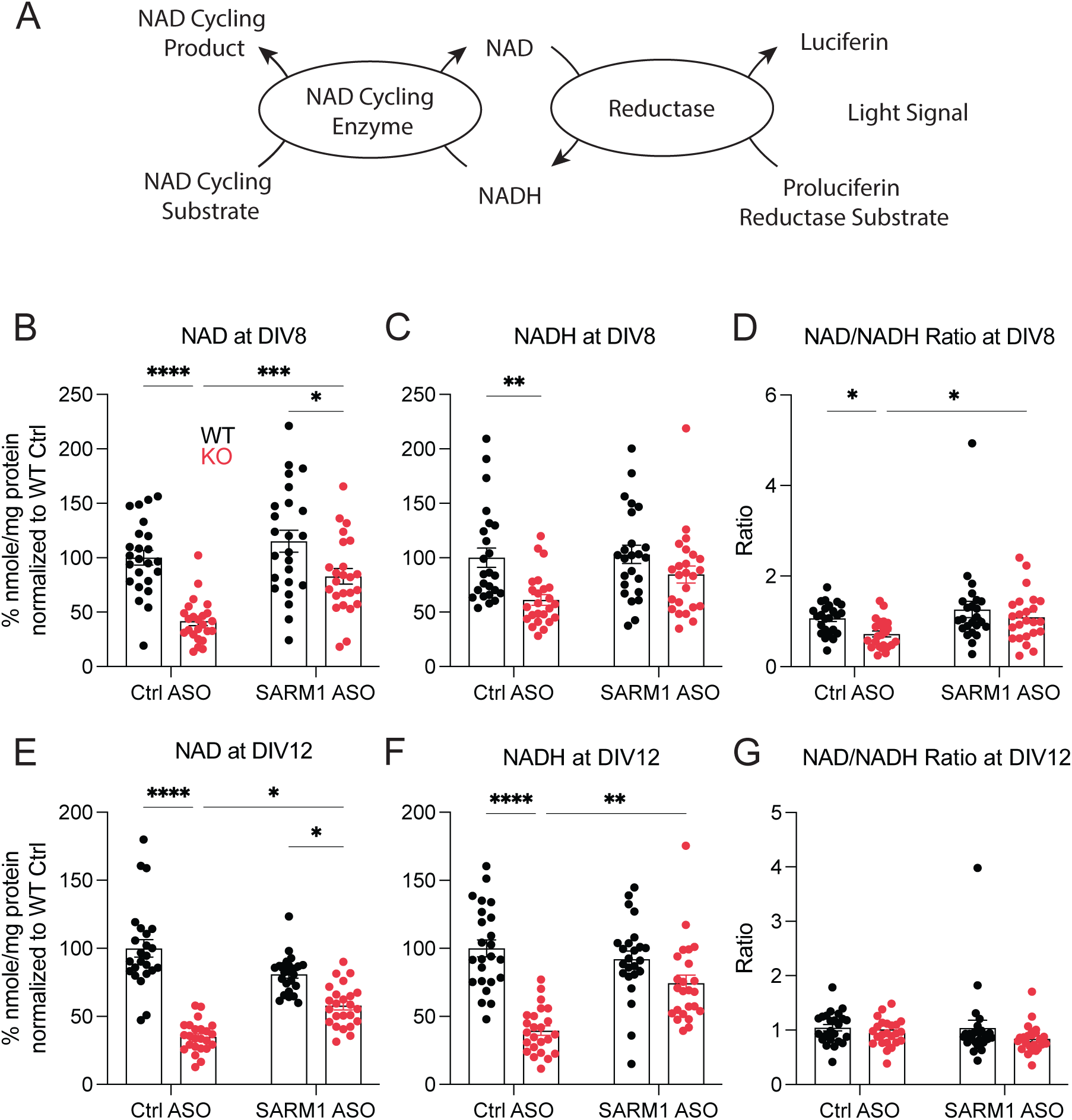
(A) Schematic representation of the reactions involved in the NAD^+^/NADH-Glo™ bioluminescent assay (B) NAD quantification following DIV1-8 of ASO treatment. Data was analyzed using two-way ANOVA with Tukey’s multiple comparisons test. *, p=0.0136, ***, p=0.0009, ****, p<0.001. (C) NADH quantification following DIV1-8 of ASO treatment. Data was analyzed using Kruskal-Wallis test with Dunn’s multiple comparisons test. **, p=0.002. (D) NAD/NADH ratio quantification following DIV1-8 of ASO treatment. Data was analyzed using Kruskal-Wallis test with Dunn’s multiple comparisons test. WT Ctrl ASO vs. KO Ctrl ASO *, p=0.0139, KO Ctrl ASO vs KO SARM1 ASO *, p=0.0334. n=24 WT Ctrl ASO, 24 WT SARM1 ASO, 24 KO Ctrl ASO, 24 KO SARM1 ASO. (E) NAD quantification following DIV1-12 of ASO treatment. Data was analyzed using Kruskal-Wallis test with Dunn’s multiple comparisons test. KO Ctrl ASO vs KO SARM1 ASO *, p=0.0182, WT SARM1 ASO vs KO SARM1 ASO *, p=0.0127, ****, p<0.001. (F) NADH quantification following DIV1-12 of ASO treatment. Data was analyzed using Kruskal-Wallis test with Dunn’s multiple comparisons test. **, p=0.0027, ****, p<0.001. (G) NAD/NADH ratio quantification following DIV1-12 of ASO treatment. Data was analyzed using Kruskal-Wallis test with Dunn’s multiple comparisons test. n=24 WT Ctrl ASO, 24 WT SARM1 ASO, 24 KO Ctrl ASO, 24 KO SARM1 ASO. Samples were collected from 3 independent experiments. All bar graphs represent mean ± SEM.

### Metabolic functional analyses show the importance of the NMNAT2-SARM1 axis in neuronal glucose metabolism

The NAD^+^/NADH redox potential is essential to drive glycolysis and oxidative phosphorylation. NAD⁺/NADH redox potentials dictate the rate of glycolytic flux, serve as a vital electron acceptor, and are essential for maintaining glucose-derived ATP production (51). To determine how NAD^+^ reduction in neurons impacts energy homeostasis, we first determined if glycolysis is impaired in KO neurons and whether SARM1 KD or exogenous NAD supplementation rescues it. Glycolytic Rate Seahorse assay was conducted with DIV8 and DIV12 WT and KO neurons. Glycolytic Rate assay measures proton efflux rate (PER) at basal, induced, and compensatory stages, which is then transformed into glycoPER to assess cellular glycolytic capacity. First, we injected NMDA and glycine to induce chemical long-term potentiation (LTP) (52) and measured glycolysis under enhanced synaptic transmission (Figure 4A). Next, we measured compensatory responses by injecting a combination of rotenone and antimycin to block mitochondrial respiration and push the cells to their maximum glycolytic capacity. Finally, 2DG was injected to block glycolysis to confirm that the previous measurements were primarily derived from glycolysis (53) (Figure 4B).

**Figure 4.**
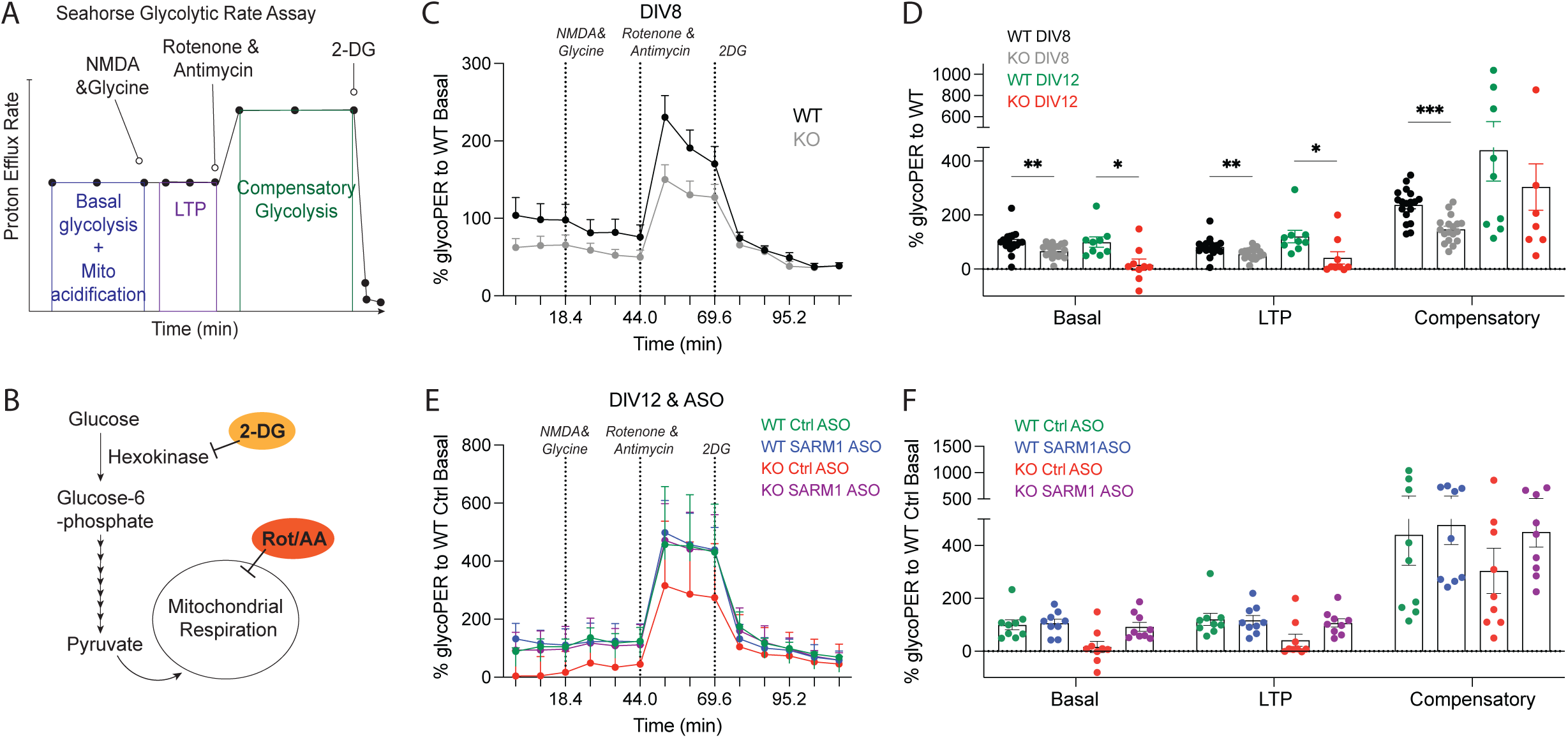
(A) Seahorse Induced Glycolytic Rate test profile. (B) Illustration of the glycolysis pathway and the targets for rotenone & antimycin, and 2DG. (C) Glycolytic rate assay response curve of NMNAT2 WT and KO neurons at DIV8. (D) Normalized glycoPER rates of DIV8 and DIV12 WT and KO neurons at basal, LTP, and compensatory steps. Data was analyzed using Mann-Whitney test and Holm-Šidák’s multiple comparisons test. DIV8 WT vs KO Basal; **, p= 0.0026. DIV8 WT vs KO LTP; **, p= 0.0021. DIV8 WT vs KO Compensatory; ****, p<0.001. DIV12 WT vs KO Basal; *, p=0.0106. DIV12 WT vs KO LTP; *, p=0.0142. n= 18 WT DIV8, 18 KO DIV8, 9 WT DIV12, 9 KO DIV12. DIV8 samples were collected from 4 independent batches and DIV12 samples were collected from 2 independent batches. (E) Glycolytic rate assay response curve of NMNAT2 WT and KO neurons treated with Ctrl or SARM1 ASO at DIV12. (F) Normalized glycoPER rates of DIV12 WT and KO neurons treated with ASO at basal, LTP, and compensatory steps. Data was analyzed using Kruskal-Wallis test with Dunn’s multiple comparisons test. *, p= 0.0459. n= 9 WT Ctrl ASO, 9 WT SARM1 ASO, 9 KO Ctrl ASO, 9 KO SARM1 ASO. Samples collected from 2 independent batches. All bar graphs represent mean ± SEM.

We observed significantly reduced glycoPER responses in NMNAT2 KO neurons compared to WT through basal, LTP, and compensatory phases at DIV8 (Figure 4C-D). Similar reductions were observed with DIV12 KO neurons, except for compensatory responses. We next treated neurons with SARM1 ASO to determine if reducing SARM1 could restore glycolytic capacity. We found that treatment with SARM1 ASO KO restored glycoPER responses in all three phases to control levels, whereas Ctrl ASO treatment exerted no rescue (Figure 4E-F).

SARM1 has been shown to localize in mitochondria (54, 55). To determine if the NMNAT2-SARM1 axis mitochondrial function is impaired in NMNAT2 KO neurons, and whether SARM1 KD or NAD^+^ supplementation can restore its capacity, the Mito Stress Seahorse assay was employed to measure OCR at basal and maximal respiration conditions, and ATP production capacity (Figure 5A). The injection paradigm begins with oligomycin, which inhibits complex V of the electron transport chain, and the resulting drop in OCR provides information on the amount of ATP being produced by the mitochondria. This is followed by an injection of FCCP, a protonophore that uncouples the mitochondria’s proton gradient and stimulates rapid oxygen consumption to calculate maximal respiratory capacity in the mitochondria. Finally, a combination of rotenone and antimycin is injected to block complexes I and III of the electron transport chain and completely shut down the mitochondria, revealing OCR readings from proton leak (56) (Figure 5B).

**Figure 5.**
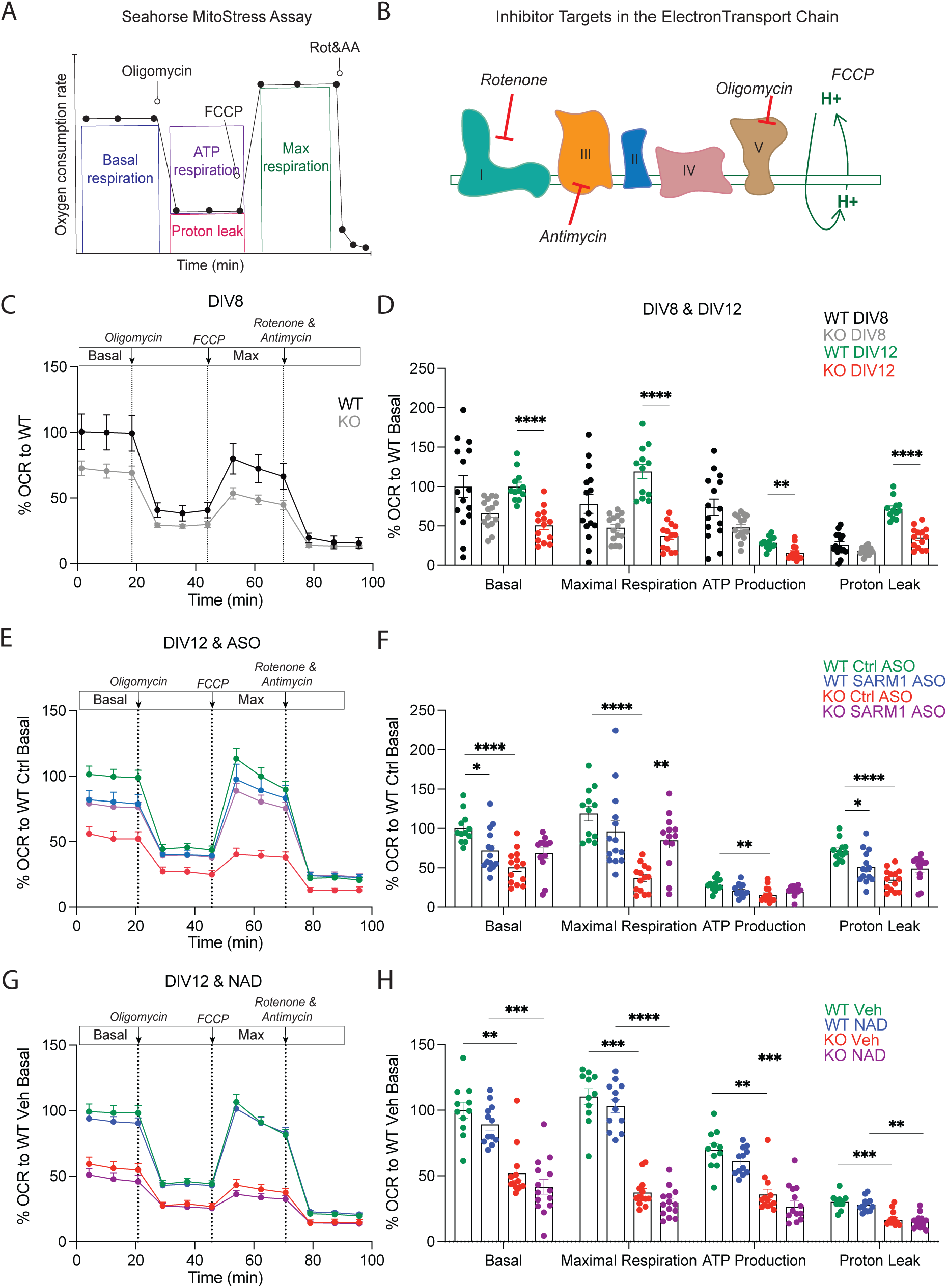
(A) Seahorse MitoStress Assay test profile. (B) Mitochondria respiratory chain complex and illustration of targets for rotenone, antimycin, oligomycin and FCCP. (C) MitoStress assay response curve of NMNAT2 WT and KO neurons at DIV 8. (D) Normalized OCR rates of DIV8 and DIV12 WT and KO neurons at basal, maximal respiration, ATP production, and proton leak steps. Data was analyzed using Mann-Whitney test and Holm-Šidák’s multiple comparisons test. **, p= 0.0013, ****, p<0.001. n= 15 WT DIV8, 15 KO DIV8, 12 WT DIV12, 14 KO DIV12. (E) MitoStress assay response curve of NMNAT2 WT and KO neurons treated with Ctrl or SARM1 ASO at DIV12. (F) Normalized OCR rates of DIV12 WT and KO neurons treated with ASO at basal, maximal, ATP production, and proton leak steps. Data was analyzed using Kruskal-Wallis test with Dunn’s multiple comparisons test. Basal; *, p= 0.0238, ****, p<0.001. Maximal respiration; **, p= 0.0045, ****, p<0.001. ATP production; **, p= 0.0022. Proton leak; *, p=0.032, ****, p<0.0001. n= 12 WT Ctrl ASO, 14 WT SARM1 ASO, 14 KO Ctrl ASO, 14 KO SARM1 ASO. (G) MitoStress assay response curve of NMNAT2 WT and KO neurons treated with vehicle or NAD at DIV12. (H) Normalized OCR rates of DIV12 WT and KO neurons treated with vehicle or NAD at basal, maximal respiration, ATP production, and proton leak steps. Data was analyzed using Kruskal-Wallis test with Dunn’s multiple comparisons test. Basal; **, p= 0.0013, ***, p=0.0004. Maximal respiration; ***, p= 0.0004, ****, p<0.001. ATP production; **, p=0.0017, ***, p=0.0003. Proton leak; **, p= 0.0012, ***, p=0.0008. n= WT Veh 11, WT NAD 12, KO Veh 13, KO NAD 14. Samples collected from 3 independent batches. All bar graphs represent mean ± SEM.

Our data show that NMNAT2 KO neurons exhibit normal mitochondrial function at DIV8 (Figure 5C). However, at DIV12, OCR was significantly reduced in NMNAT2 KO neurons compared to WT controls across all test measurements (Figure 5D). SARM1 ASO treatment normalized the mitochondrial function of DIV12 KO neurons (Figure 5E-F), with similar OCR readings between SARM1-ASO-treated-NMNAT2 KO and -WT neurons for basal, maximum respiration, and ATP production phases. Ctrl ASO treatment did not rescue mitochondrial deficits in DIV12 NMNAT2 KO neurons. To our surprise, exogenous NAD^+^ supplementation did not rescue the mitochondrial function of DIV12 KO neurons (Figure 5G-H). Taken together, our data show that mitochondrial function is significantly impaired in NMNAT2 KO neurons at DIV12 in a SARM1-dependent manner.

### Temporal profiling of APP-CTF accumulations and the proteomic profiles revealed an initial progression phase followed by an accelerating phase

To determine when APP-CTFs start to accumulate and the temporal progression in relationship to glycolysis and mitochondria dysfunction onsets, we examined the abundance of APP-FL and APP-CTFs in DIV5/8/10/14 WT and NMNAT2 KO neurons with western blotting (Figure 6A). APP-FL was not different between WT and KO neurons at these timepoints (Figure 6B). At DIV5, KO neurons showed normal levels of APP-CTFs ranging from 10-18kDa (Figure 6C). By DIV8, the APP-CTFs are significantly higher in KO neurons, and such differences were accelerated between DIV12 and 14 (Figure 6C), when mitochondrial dysfunction is evident in KO neurons.

**Figure 6.**
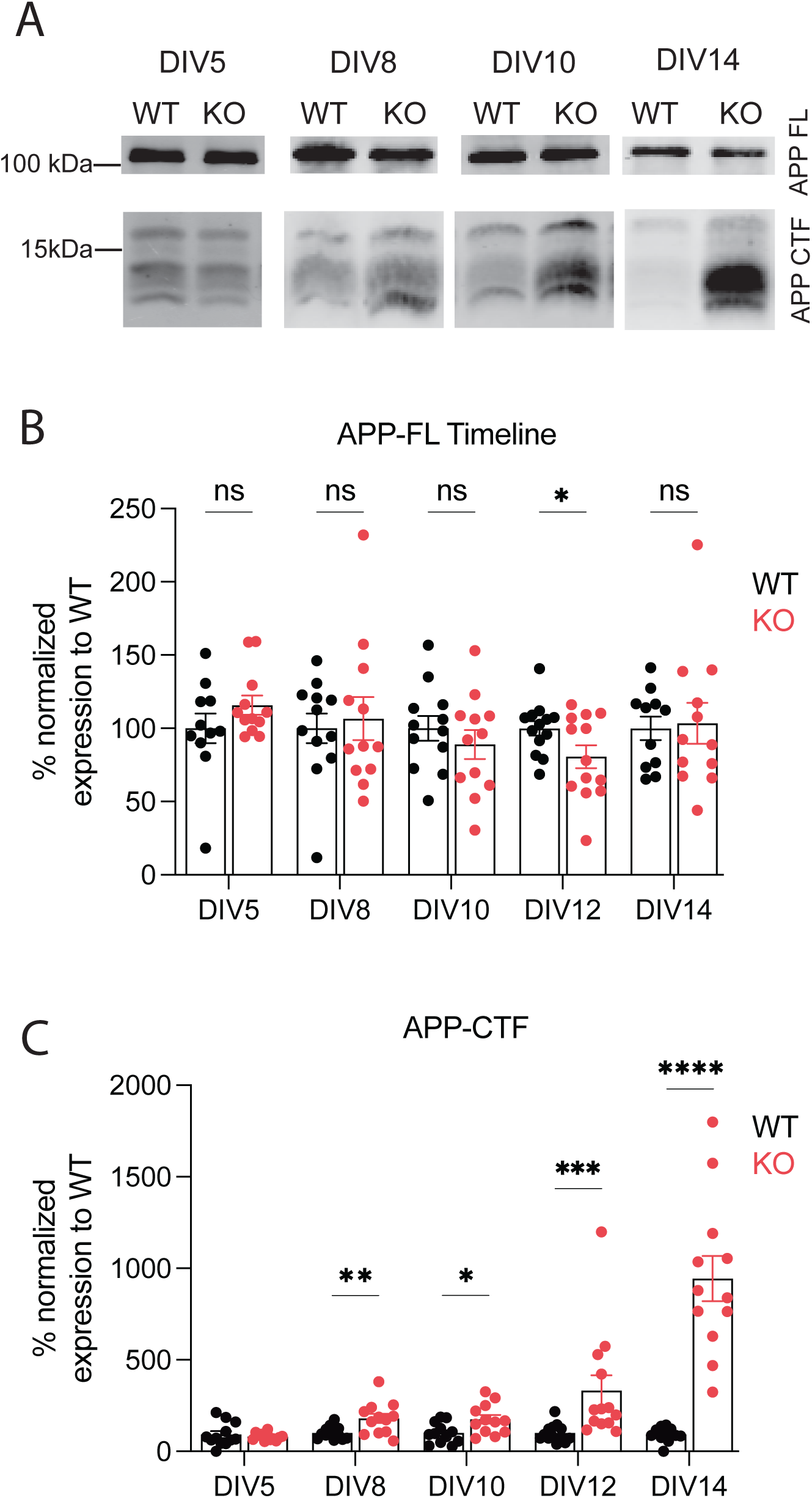
(A) Western blot of WT and KO neuronal culture lysates at DIV5, 8, 10, and 14 showing APP-FL and CTF expression. (B) Western blot analysis of APP-FL expression relative to corresponding DIV WT neurons at DIV5, 8, 10, 12 and 14. DIV5; n=11 WT, 12 KO; APP-FL was analyzed with unpaired t-test. DIV8; n=12 WT, 12 KO; APP-FL was analyzed with Mann-Whitney test. DIV10; n=12 WT, 12 KO; APP-FL was analyzed with unpaired t-test. DIV12; n=13 WT, 13 KO; APP-FL was analyzed with unpaired t-test. *, p=0.049. DIV14; n=11 WT, 12 KO; APP-FL was analyzed with Mann-Whitney test. Samples were collected from 3 independent experiments. (C) Western blot analysis of APP-CTF expression relative to corresponding DIV WT neurons at DIV5, 8, 10, 12 and 14. DIV5; n=11 WT, 12 KO; APP-CTF was analyzed with Mann-Whitney test. DIV8; n=12 WT, 12 KO; APP-CTF was analyzed with Mann-Whitney test. **, p=0.0083. DIV10; n=12 WT, 12 KO; APP-CTF was analyzed with unpaired t-test. *, p=0.015. DIV12; n=13 WT, 12 KO; APP-CTF was Mann-Whitney test. ***, p=0.0002. DIV14; n=11 WT, 12 KO; APP-CTF was analyzed with Mann-Whitney test. ****, p= <0.0001. Samples were collected from 3 independent experiments. All bar graphs represent mean ± SEM.

To explore the mechanisms affected by neuronal NAD^+^ reduction due to NMNAT2 loss, proteomic analyses were conducted on WT and NMNAT2 KO neurons at both DIV8, the stage when APP processing deficits begin to appear, and at DIV12, when APP-CTF accumulations enter the accelerated phase. Comparing KO and WT neurons at DIV8, we found only 42 significantly up-regulated and 3 significantly down-regulated proteins (Figure 7A). Among the upregulated proteins, multiple kinesin superfamily members (Kif1a, Kif1b, Kif5a, Kif5c, Kif3a, and Kif3c) were elevated, along with MAP kinase pathway components Mapk8ip1, Mapk8ip3, and Map2k2. Mapk8ip1 and Mapk8ip3 function as c-Jun N-terminal kinase (JNK) scaffold proteins that associate with kinesin motors, linking mitogen-activated protein kinases (MAPK) signaling to axonal transport machinery (57, 58) (Figure 7B). GO enrichment analysis identified significant overrepresentation of biological processes related to protein localization, endosomal and lysosomal pathways, MAPK cascade signaling, microtubule organization, and dendritic development (Figure 7C). These findings indicate coordinated upregulation of proteins involved in intracellular transport, vesicular trafficking, cytoskeletal organization, and MAPK signaling at the onset of the phenotype.

**Figure 7.**
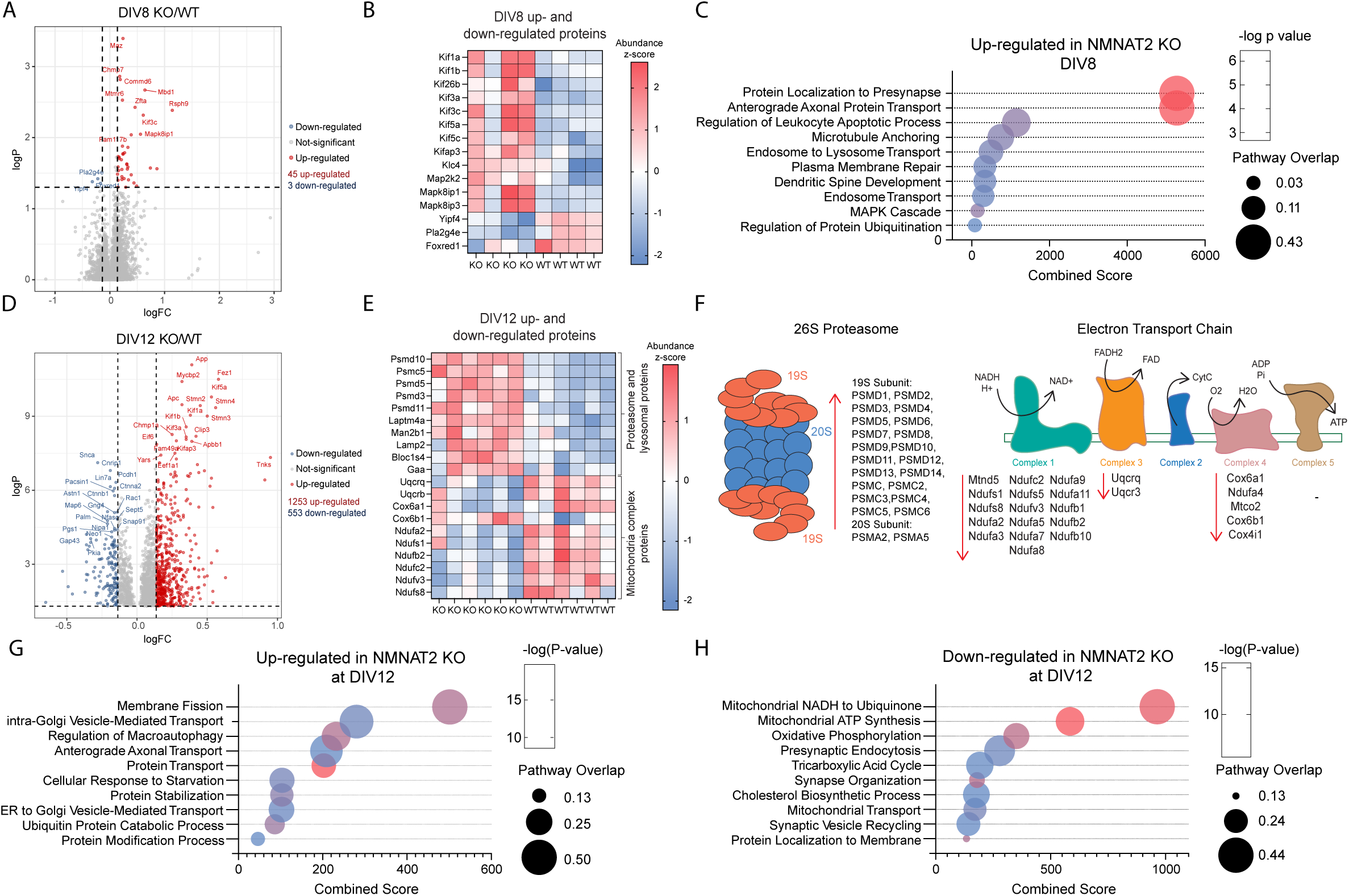
(A) Volcano plot of abundance ratio for all identified genes at DIV8 in WT and KO samples from proteomic analysis. Significantly down-regulated genes in KO samples are shown in blue (n=3), and significantly up-regulated genes are shown in red (n=42). (B) Heatmap of protein abundance across DIV8 WT and KO samples. Values were z-scored per gene, with colors indicating relative abundance compared to each protein’s mean across samples (red>0; blue<0). (C) Bubble plot of enriched GO Biological processes from up-regulated gene list at DIV8. (D) Volcano plot of abundance ratio for all identified genes at DIV12 in WT and KO samples from proteomic analysis. Significantly down-regulated genes in KO neurons are shown in blue (n=553), and significantly up-regulated proteins are shown in red (n=1253). (E) Heatmap of protein abundance across DIV8 WT and KO samples. Values were z-scored per gene, with colors indicating relative abundance compared to each protein’s mean across samples (red>0; blue<0). (F) Illustration of the 26S proteasome and its corresponding up-regulated genes in NMNAT2 KO neurons at DIV12 (left) and illustration of mitochondria complexes along with their corresponding down-regulated genes in NMNAT2 KO neurons at DIV12 (right). Arrow direction indicates down or up-regulation of genes. (G) Bubble plot of enriched GO Biological processes from up-regulated gene list at DIV12. (H) Bubble plot of enriched GO Biological processes from down-regulated gene list at DIV12. DIV8 n=4 WT, 4KO. DIV12 n= 6 WT, 6 KO.

At DIV12, the NMNAT2 KO neuron proteomic profile was vastly different from WT controls, with 1253 up-regulated proteins and 533 down-regulated proteins (Figure 7D). Examination of differentially expressed proteins revealed coordinated changes within distinct protein families and pathway-associated groups. Specifically, proteasomal subunits were consistently upregulated in NMNAT2 KO neurons, whereas several components of the mitochondria electron transport chain complexes were downregulated (Figure 7E). The 26S proteasome has 33 unique protein subunits, and 22 of these proteins were up-regulated in NMNAT2 KO neurons (Figure 7F). 16 of the 45 mitochondria complex 1 subunit proteins, along with proteins associated with complex 3 and complex 4, were downregulated in NMNAT2 KO neurons (Figure 7F). GO enrichment analysis of upregulated proteins identified processes related to protein transport, macroautophagy, cellular response to starvation, and protein ubiquitination (Figure 7G); whereas downregulated proteins were enriched for pathways associated with mitochondrial function, oxidative phosphorylation, synaptic organization, and protein localization (Figure 7H).

Protein-protein interactions analyzed with STRING further supported the above pathways (Supp. Figure 2). At DIV8, upregulated proteins clustered into a network associated with microtubule dynamics, kinesin complexes, and protein transport (Supp. Figure 2A). At DIV12, upregulated proteins formed three major clusters linked to protein translation, protein folding, and ubiquitin-mediated proteasomal degradation (Supp. Figure 2B), while downregulated proteins clustered within glycolysis, pyruvate metabolism, oxidative phosphorylation, and GTPase signaling networks (Supp. Figure 2C). Collectively, these data suggest NMNAT2 loss results in alterations in transport machinery at DIV8, while proteostasis and mitochondrial metabolic function at DIV12.

## Discussion

The NMNAT2–SARM1 axis is a key regulator of neuronal NAD⁺ homeostasis and energy metabolism (25). Here, we show that neuronal NMNAT2 loss induces SARM1-dependent accumulation of APP-CTF. Temporal profiling shows that APP-CTF accumulation in NMNAT2 KO neurons coincides with early reductions in NAD+/NADH redox potential and impaired glycolysis, while mitochondrial function is initially preserved but declines at later stages. This delayed mitochondrial dysfunction is accompanied by extensive proteomic remodeling and accelerated APP-CTF accumulation. Notably, SARM1 KD restores APP processing, glycolysis, and oxidative phosphorylation in NMNAT2 KO neurons, whereas NAD⁺ supplementation provides only partial rescue. Together, these findings support a model in which disruption of NAD^+^-dependent redox homeostasis drives defects in APP processing, positioning the NMNAT2–SARM1 axis as a central regulator linking neuronal bioenergetics to proteostasis.

### Progressive Metabolic Failure Disrupts APP Processing

APP-CTFs begin to accumulate in NMNAT2 KO neurons at DIV8 (Figure 6), coinciding with previously reported axonal transport deficits in APP-containing cargos (25). This timepoint corresponds to a stage of active synaptogenesis and increasing energetic demand in neurons (59, 60). The temporal alignment between rising energy demand, reduced NAD⁺ levels (Figure 3), and APP-CTF accumulation suggests that impaired energy homeostasis is a primary cause of defective APP processing.

Metabolic profiling revealed significant glycolytic deficits at DIV8 (Figure 4), consistent with prior observations in NMNAT2-deficient axons (25). In contrast, mitochondrial oxidative phosphorylation deficits emerge later at DIV12 (Figure 5), coinciding with a sharp increase in APP-CTF levels (Figure 6C). Proteomic analysis further shows downregulation of mitochondrial complex I proteins in NMNAT2 KO neurons at DIV12 (Figure 7F), but not at DIV8, raising the possibility that progressive mitochondrial dysfunction amplifies defects in APP processing.

Mitochondrial dysfunction is linked to impaired mitochondrial protein synthesis and disrupted organelle maintenance (61). This is consistent with the coordinated reduction in respiratory complex components observed in our proteomic data. In addition, mitochondrial dysfunction and abnormal APP processing can influence each other (62), potentially establishing a pathogenic feedback loop that exacerbates proteostatic stress and contributes to the accelerated accumulation of APP-CTFs after DIV12.

### APP Trafficking Defects Underlie Abnormal Processing in NMNAT2 KO Neuron

Proteomic profiling at DIV8 revealed upregulation of kinesin motor proteins and MAPK pathway components, including JNK scaffold proteins. JNK signaling, which is responsive to metabolic stress and regulates kinesin activity (63, 64), may contribute to altered transport dynamics. Early APP axonal transport deficits observed in NMNAT2 KO axons (25) suggest that increased levels of axonal transport components may represent a compensatory response. However, altered motor protein expression and stress signaling could also impair trafficking through secretory and endocytic pathways, promoting APP retention in secretase-rich compartments and enhancing its cleavage (65, 66). Disrupted endosomal and trans-Golgi-network trafficking may further alter APP processing (67, 68).

At later stages (DIV12), proteomic analysis revealed upregulation of unfolded protein response (UPR)-related proteins, including increased proteasome components and ubiquitin-mediated protein catabolism (69, 70), consistent with ER stress (71). These findings support a model in which NMNAT2 loss disrupts ER-to-Golgi trafficking and proteostatic balance. Additionally, the increase of APP-CTFs themselves may contribute to cellular stress, as CTF-99 fragments can impair lysosomal function, disrupt APP transport (72, 73), and promote neuronal hyperexcitability (34). Together, these results suggest that NMNAT2 loss leads to defects in protein trafficking and proteostasis that contribute to abnormal APP processing.

### SARM1 Amplifies Metabolic and Proteostatic Dysfunction after NMNAT2 Loss

SARM1 activation caused by NMNAT2 loss amplifies metabolic failure through NAD⁺ consumption, contributing to downstream mitochondrial dysfunction (74, 75). SARM1 KD restores APP-CTF levels (Figure 2A-C), glycolytic function (Figure 4E-F), and mitochondrial oxidative capacity (Figure 5E-F), reversing both bioenergetic and proteostatic defects in NMNAT2 KO neurons despite incomplete normalization of NAD⁺ levels. This partial recovery is consistent with the continued absence of NMNAT2, the major NAD⁺ synthesizing enzyme in neurons. SARM1 KD prevents further pathological NAD⁺ consumption but does not replenish NAD⁺ pools. Our previous work shows NAD⁺ supplementation rescues axonal glycolysis (25). Here, we found NAD⁺ partially rescued APP-CTF levels but failed to restore mitochondrial respiration, potentially reflecting compartmental limitations or incomplete correction of intracellular redox balance. These findings suggest that SARM1-mediated pathology extends beyond NAD⁺ depletion alone and involves a broader disruption of metabolic and redox homeostasis. The effective rescue by SARM1 KD demonstrates that blocking this amplification step enables neurons to endure suboptimal energy conditions.

## Conclusion

In summary, loss of NMNAT2 triggers a progressive, SARM1-dependent disruption of neuronal redox homeostasis, energy metabolism, and proteostasis, leading to accumulation of APP-CTFs. These findings identify the NMNAT2–SARM1 axis as a central regulatory pathway linking NAD^+^-dependent redox balance to protein processing and identify SARM1 as a key mediator that translates metabolic stress into sustained dysfunction. This work emphasizes metabolic dysfunction as a driver of neuronal pathology and supports SARM1 inhibition as a promising therapeutic approach. Future studies should determine whether enhancing mitochondrial function or redox balance can prevent proteinopathy and whether SARM1 is required for activation of stress pathways such as UPR and proteasome system.

## Acknowledgments

This work was supported by the National Institutes of Health grant NINDS NS086794 (HCL). We thank Zhen-Xian Niou, Jui-Yen Huang, Nino A. Espinas, and Scott Barton for their technical support.

## REFERENCES

1. Yu L, Jin J, Xu Y, Zhu X. Aberrant Energy Metabolism in Alzheimer’s Disease. J Transl Int Med. 2022;10(3):197–206.

2. Mosconi L, Pupi A, De Leon MJ. Brain glucose hypometabolism and oxidative stress in preclinical Alzheimer’s disease. Ann N Y Acad Sci. 2008;1147:180–95.

3. Watanabe H, Shima S, Kawabata K, Mizutani Y, Ueda A, Ito M. Brain network and energy imbalance in Parkinson’s disease: linking ATP reduction and alpha-synuclein pathology. Front Mol Neurosci. 2024;17:1507033.

4. Mochel F, Haller RG. Energy deficit in Huntington disease: why it matters. J Clin Invest. 2011;121(2):493–9.

5. Ganguly G, Chakrabarti S, Chatterjee U, Saso L. Proteinopathy, oxidative stress and mitochondrial dysfunction: cross talk in Alzheimer’s disease and Parkinson’s disease. Drug Des Devel Ther. 2017;11:797–810.

6. Pievani M, Filippini N, van den Heuvel MP, Cappa SF, Frisoni GB. Brain connectivity in neurodegenerative diseases--from phenotype to proteinopathy. Nat Rev Neurol. 2014;10(11):620–33.

7. Braak H, Braak E. Neuropathological stageing of Alzheimer-related changes. Acta Neuropathol. 1991;82(4):239–59.

8. Camandola S, Mattson MP. Brain metabolism in health, aging, and neurodegeneration. EMBO J. 2017;36(11):1474–92.

9. Yin F, Sancheti H, Patil I, Cadenas E. Energy metabolism and inflammation in brain aging and Alzheimer’s disease. Free Radic Biol Med. 2016;100:108–22.

10. Pathak D, Berthet A, Nakamura K. Energy failure: does it contribute to neurodegeneration? Ann Neurol. 2013;74(4):506–16.

11. Na D, Zhang Z, Meng M, Li M, Gao J, Kong J, et al. Energy Metabolism and Brain Aging: Strategies to Delay Neuronal Degeneration. Cell Mol Neurobiol. 2025;45(1):38.

12. Sheline YI, Raichle ME. Resting state functional connectivity in preclinical Alzheimer’s disease. Biol Psychiatry. 2013;74(5):340–7.

13. Yang Y, Sauve AA. NAD(+) metabolism: Bioenergetics, signaling and manipulation for therapy. Biochim Biophys Acta. 2016;1864(12):1787–800.

14. Schiuma G, Lara D, Clement J, Narducci M, Rizzo R. Nicotinamide Adenine Dinucleotide: The Redox Sensor in Aging-Related Disorders. Antioxid Redox Signal. 2024.

15. Wilson N, Kataura T, Korsgen ME, Sun C, Sarkar S, Korolchuk VI. The autophagy-NAD axis in longevity and disease. Trends Cell Biol. 2023;33(9):788–802.

16. Buttgereit F, Brand MD. A hierarchy of ATP-consuming processes in mammalian cells. Biochem J. 1995;312 (Pt 1)(Pt 1):163–7.

17. Liu YJ, Chern Y. Contribution of Energy Dysfunction to Impaired Protein Translation in Neurodegenerative Diseases. Front Cell Neurosci. 2021;15:668500.

18. Ottens F, Franz A, Hoppe T. Build-UPS and break-downs: metabolism impacts on proteostasis and aging. Cell Death Differ. 2021;28(2):505–21.

19. Schroeder HT, De Lemos Muller CH, Heck TG, Krause M, Homem de Bittencourt PI, Jr. The dance of proteostasis and metabolism: Unveiling the caloristatic controlling switch. Cell Stress Chaperones. 2024;29(1):175–200.

20. Berger F, Lau C, Dahlmann M, Ziegler M. Subcellular compartmentation and differential catalytic properties of the three human nicotinamide mononucleotide adenylyltransferase isoforms. J Biol Chem. 2005;280(43):36334–41.

21. Raffaelli N, Sorci L, Amici A, Emanuelli M, Mazzola F, Magni G. Identification of a novel human nicotinamide mononucleotide adenylyltransferase. Biochem Biophys Res Commun. 2002;297(4):835–40.

22. Yan T, Feng Y, Zheng J, Ge X, Zhang Y, Wu D, et al. Nmnat2 delays axon degeneration in superior cervical ganglia dependent on its NAD synthesis activity. Neurochem Int. 2010;56(1):101–6.

23. Ali YO, Allen HM, Yu L, Li-Kroeger D, Bakhshizadehmahmoudi D, Hatcher A, et al. NMNAT2:HSP90 Complex Mediates Proteostasis in Proteinopathies. PLoS Biol. 2016;14(6):e1002472.

24. Harlan BA, Killoy KM, Pehar M, Liu L, Auwerx J, Vargas MR. Evaluation of the NAD(+) biosynthetic pathway in ALS patients and effect of modulating NAD(+) levels in hSOD1-linked ALS mouse models. Exp Neurol. 2020;327:113219.

25. Yang S, Niou ZX, Enriquez A, LaMar J, Huang JY, Ling K, et al. NMNAT2 supports vesicular glycolysis via NAD homeostasis to fuel fast axonal transport. Mol Neurodegener. 2024;19(1):13.

26. Niou ZX, Yang S, Enriquez A, Espinas NA, Sri A, Hines CD, et al. NAD(+) Reduction in Glutamatergic Neurons Induces Lipid Catabolism and Neuroinflammation in the Brain via SARM1. Adv Sci (Weinh). 2025:e09950.

27. Figley MD, Gu W, Nanson JD, Shi Y, Sasaki Y, Cunnea K, et al. SARM1 is a metabolic sensor activated by an increased NMN/NAD(+) ratio to trigger axon degeneration. Neuron. 2021;109(7):1118–36 e11.

28. Gerdts J, Brace EJ, Sasaki Y, DiAntonio A, Milbrandt J. SARM1 activation triggers axon degeneration locally via NAD(+) destruction. Science. 2015;348(6233):453–7.

29. Chow VW, Mattson MP, Wong PC, Gleichmann M. An overview of APP processing enzymes and products. Neuromolecular Med. 2010;12(1):1–12.

30. O’Brien RJ, Wong PC. Amyloid precursor protein processing and Alzheimer’s disease. Annu Rev Neurosci. 2011;34:185–204.

31. Vaillant-Beuchot L, Mary A, Pardossi-Piquard R, Bourgeois A, Lauritzen I, Eysert F, et al. Accumulation of amyloid precursor protein C-terminal fragments triggers mitochondrial structure, function, and mitophagy defects in Alzheimer’s disease models and human brains. Acta Neuropathol. 2021;141(1):39–65.

32. Vaillant-Beuchot L, Eysert F, Duval B, Kinoshita PF, Pardossi-Piquard R, Bauer C, et al. The amyloid precursor protein and its derived fragments concomitantly contribute to the alterations of mitochondrial transport machinery in Alzheimer’s disease. Cell Death Dis. 2024;15(5):367.

33. Pera M, Larrea D, Guardia-Laguarta C, Montesinos J, Velasco KR, Agrawal RR, et al. Increased localization of APP-C99 in mitochondria-associated ER membranes causes mitochondrial dysfunction in Alzheimer disease. EMBO J. 2017;36(22):3356–71.

34. Kapadia A, Schuhmann F, Daskin E, Walter J, Lindahl I, Rahmani N, et al. Amyloidogenic proteolysis of APP regulates glutamatergic presynaptic function. bioRxiv. 2025:2025.08.01.667924.

35. Luo M, Zhou J, Sun C, Chen W, Fu C, Si C, et al. APP beta-CTF triggers cell-autonomous synaptic toxicity independent of Abeta. Elife. 2025;13.

36. Lauritzen I, Pardossi-Piquard R, Bourgeois A, Pagnotta S, Biferi MG, Barkats M, et al. Intraneuronal aggregation of the beta-CTF fragment of APP (C99) induces Abeta-independent lysosomal-autophagic pathology. Acta Neuropathol. 2016;132(2):257–76.

37. Bretou M, Sannerud R, Escamilla-Ayala A, Leroy T, Vrancx C, Van Acker ZP, et al. Accumulation of APP C-terminal fragments causes endolysosomal dysfunction through the dysregulation of late endosome to lysosome-ER contact sites. Dev Cell. 2024;59(12):1571–92 e9.

38. Hung COY, Livesey FJ. Altered gamma-Secretase Processing of APP Disrupts Lysosome and Autophagosome Function in Monogenic Alzheimer’s Disease. Cell Rep. 2018;25(13):3647–60 e2.

39. Miranda AM, Lasiecka ZM, Xu Y, Neufeld J, Shahriar S, Simoes S, et al. Neuronal lysosomal dysfunction releases exosomes harboring APP C-terminal fragments and unique lipid signatures. Nat Commun. 2018;9(1):291.

40. Eggert S, Thomas C, Kins S, Hermey G. Trafficking in Alzheimer’s Disease: Modulation of APP Transport and Processing by the Transmembrane Proteins LRP1, SorLA, SorCS1c, Sortilin, and Calsyntenin. Mol Neurobiol. 2018;55(7):5809–29.

41. Tan JZA, Gleeson PA. The role of membrane trafficking in the processing of amyloid precursor protein and production of amyloid peptides in Alzheimer’s disease. Biochim Biophys Acta Biomembr. 2019;1861(4):697–712.

42. Yu Y, Zhou RZ, Nilsson P, Winblad B, Tjernberg LO, Schedin-Weiss S. Altered APP trafficking drives amyloidogenic processing in primary neurons from the App(NL-F) knock-in mouse model of Alzheimer’s disease. Neurobiol Dis. 2025;216:107129.

43. Wilkins HM, Swerdlow RH. Amyloid precursor protein processing and bioenergetics. Brain Res Bull. 2017;133:71–9.

44. Yang TT, Shih YS, Chen YW, Kuo YM, Lee CW. Glucose regulates amyloid beta production via AMPK. J Neural Transm (Vienna). 2015;122(10):1381–90.

45. Cai Z, Yan LJ, Li K, Quazi SH, Zhao B. Roles of AMP-activated protein kinase in Alzheimer’s disease. Neuromolecular Med. 2012;14(1):1–14.

46. Hicks AN, Lorenzetti D, Gilley J, Lu B, Andersson KE, Miligan C, et al. Nicotinamide mononucleotide adenylyltransferase 2 (Nmnat2) regulates axon integrity in the mouse embryo. PLoS One. 2012;7(10):e47869.

47. Schindelin J, Arganda-Carreras I, Frise E, Kaynig V, Longair M, Pietzsch T, et al. Fiji: an open-source platform for biological-image analysis. Nat Methods. 2012;9(7):676–82.

48. Stachel SJ, Coburn CA, Steele TG, Jones KG, Loutzenhiser EF, Gregro AR, et al. Structure-based design of potent and selective cell-permeable inhibitors of human beta-secretase (BACE-1). J Med Chem. 2004;47(26):6447–50.

49. Czvitkovich S, Duller S, Mathiesen E, Lorenzoni K, Imbimbo BP, Hutter-Paier B, et al. Comparison of pharmacological modulation of APP metabolism in primary chicken telencephalic neurons and in a human neuroglioma cell line. J Mol Neurosci. 2011;43(3):257–67.

50. Miao X, Wu Q, Du S, Xiang L, Zhou S, Zhu J, et al. SARM1 Promotes Neurodegeneration and Memory Impairment in Mouse Models of Alzheimer’s Disease. Aging Dis. 2024;15(1):390–407.

51. Xiao W, Wang RS, Handy DE, Loscalzo J. NAD(H) and NADP(H) Redox Couples and Cellular Energy Metabolism. Antioxid Redox Signal. 2018;28(3):251–72.

52. Lu W, Man H, Ju W, Trimble WS, MacDonald JF, Wang YT. Activation of synaptic NMDA receptors induces membrane insertion of new AMPA receptors and LTP in cultured hippocampal neurons. Neuron. 2001;29(1):243–54.

53. Agliano F, Menoret A, Vella AT. Evaluating the glycolytic potential of mouse costimulated effector CD8(+) T cells ex vivo. STAR Protoc. 2022;3(2):101441.

54. Panneerselvam P, Singh LP, Ho B, Chen J, Ding JL. Targeting of pro-apoptotic TLR adaptor SARM to mitochondria: definition of the critical region and residues in the signal sequence. Biochem J. 2012;442(2):263–71.

55. Kim Y, Zhou P, Qian L, Chuang JZ, Lee J, Li C, et al. MyD88-5 links mitochondria, microtubules, and JNK3 in neurons and regulates neuronal survival. J Exp Med. 2007;204(9):2063–74.

56. Gu X, Ma Y, Liu Y, Wan Q. Measurement of mitochondrial respiration in adherent cells by Seahorse XF96 Cell Mito Stress Test. STAR Protoc. 2021;2(1):100245.

57. Verhey KJ, Rapoport TA. Kinesin carries the signal. Trends Biochem Sci. 2001;26(9):545–50.

58. Verhey KJ, Meyer D, Deehan R, Blenis J, Schnapp BJ, Rapoport TA, et al. Cargo of kinesin identified as JIP scaffolding proteins and associated signaling molecules. J Cell Biol. 2001;152(5):959–70.

59. Verstraelen P, Van Dyck M, Verschuuren M, Kashikar ND, Nuydens R, Timmermans JP, et al. Image-Based Profiling of Synaptic Connectivity in Primary Neuronal Cell Culture. Front Neurosci. 2018;12:389.

60. Grabrucker A, Vaida B, Bockmann J, Boeckers TM. Synaptogenesis of hippocampal neurons in primary cell culture. Cell Tissue Res. 2009;338(3):333–41.

61. Lu B, Guo S. Mechanisms Linking Mitochondrial Dysfunction and Proteostasis Failure. Trends Cell Biol. 2020;30(4):317–28.

62. Wilkins HM. Interactions between amyloid, amyloid precursor protein, and mitochondria. Biochem Soc Trans. 2023;51(1):173–82.

63. Johnson GL, Nakamura K. The c-jun kinase/stress-activated pathway: regulation, function and role in human disease. Biochim Biophys Acta. 2007;1773(8):1341–8.

64. de Los Reyes Corrales T, Losada-Perez M, Casas-Tinto S. JNK Pathway in CNS Pathologies. Int J Mol Sci. 2021;22(8).

65. Zhang X, Song W. The role of APP and BACE1 trafficking in APP processing and amyloid-beta generation. Alzheimers Res Ther. 2013;5(5):46.

66. Zhuravleva V, Vaz-Silva J, Zhu M, Gomes P, Silva JM, Sousa N, et al. Rab35 and glucocorticoids regulate APP and BACE1 trafficking to modulate Abeta production. Cell Death Dis. 2021;12(12):1137.

67. Haass C, Kaether C, Thinakaran G, Sisodia S. Trafficking and proteolytic processing of APP. Cold Spring Harb Perspect Med. 2012;2(5):a006270.

68. Andersen OM, Reiche J, Schmidt V, Gotthardt M, Spoelgen R, Behlke J, et al. Neuronal sorting protein-related receptor sorLA/LR11 regulates processing of the amyloid precursor protein. Proc Natl Acad Sci U S A. 2005;102(38):13461–6.

69. Chen X, Shi C, He M, Xiong S, Xia X. Endoplasmic reticulum stress: molecular mechanism and therapeutic targets. Signal Transduct Target Ther. 2023;8(1):352.

70. Win S, Than TA, Fernandez-Checa JC, Kaplowitz N. JNK interaction with Sab mediates ER stress induced inhibition of mitochondrial respiration and cell death. Cell Death Dis. 2014;5(1):e989.

71. Zhang K. Endoplasmic reticulum stress response and transcriptional reprogramming. Front Genet. 2014;5:460.

72. Lauritzen I, Pardossi-Piquard R, Bourgeois A, Pagnotta S, Biferi MG, Barkats M, et al. Correction to: Intraneuronal aggregation of the beta–CTF fragment of APP (C99) induces Abeta–independent lysosomal-autophagic pathology. Acta Neuropathol. 2023;146(4):659–60.

73. Rodrigues EM, Weissmiller AM, Goldstein LS. Enhanced beta-secretase processing alters APP axonal transport and leads to axonal defects. Hum Mol Genet. 2012;21(21):4587–601.

74. Loreto A, Cramb KML, McDermott LA, Antoniou C, Cirilli I, Caiazza MC, et al. SARM1 activation induces reversible mitochondrial dysfunction and can be prevented in human neurons by antisense oligonucleotides. Neurobiol Dis. 2025;213:106986.

75. Ko KW, Devault L, Sasaki Y, Milbrandt J, DiAntonio A. Live imaging reveals the cellular events downstream of SARM1 activation. Elife. 2021;10.

