## Supplementary figures for "NMNAT2-SARM1 Axis Drives Redox Failure and Disrupts APP Processing in Neurons"

**Supplementary Figure 1**. (A) Treatment timeline and western blot of DIV10-13 control (WT and KO) and beta-secretase inhibitor-treated neurons (WT+B-inh and KO+B-inh) showing APP-FL and CTF expression. (B) Quantification of western blot expression for APP-FL, (C) CTF cluster of bands, (D) CTF99 band, and (E) CTF83 band. n=9 WT, 9 WT+B-inh, 9 KO, 9 KO+B-inh. APP-FL and CTF data was analyzed using one-way ANOVA with Tukey's multiple comparisons test. APP-FL WT Veh vs. WT B-inh **, p=0.0058. APP-FL KO Veh vs. KO B-inh **, p=0.0048. APP-CTF WT Veh vs. KO Veh **, p=0.0024. APP-CTF WT B-inh vs. KO B-Inh ***, p=0.0001. APP-CTF99 and CTF83 data was analyzed using Mann-Whitney test. APP-CTF99 WT Veh vs. KO Veh **, p=0.004. APP-CTF83 WT Veh vs. KO Veh **, p=0.0012. APP-CTF83 WT B-inh vs. KO B-inh ****, p=<0.001. Samples were collected from 3 independent experiments. All bar graphs represent mean ± SEM.

***Supplementary Figure 2.*** (A) STRING cluster of up-regulated pathways in KO neurons at DIV8. (B) STRING clusters of up-regulated pathways in KO neurons at DIV12 (C) STRING clusters of down-regulated pathways in KO neurons at DIV12.Cytoscape's STRING functional enrichment was performed to identify cluster pathways in all clusters, and FDR value was used to pick top terms for GO cellular component, molecular function, and/or biological process. DIV8 n=4 WT, 4KO. DIV12 n= 6 WT, 6 KO.

*Supplementary Figure 1*

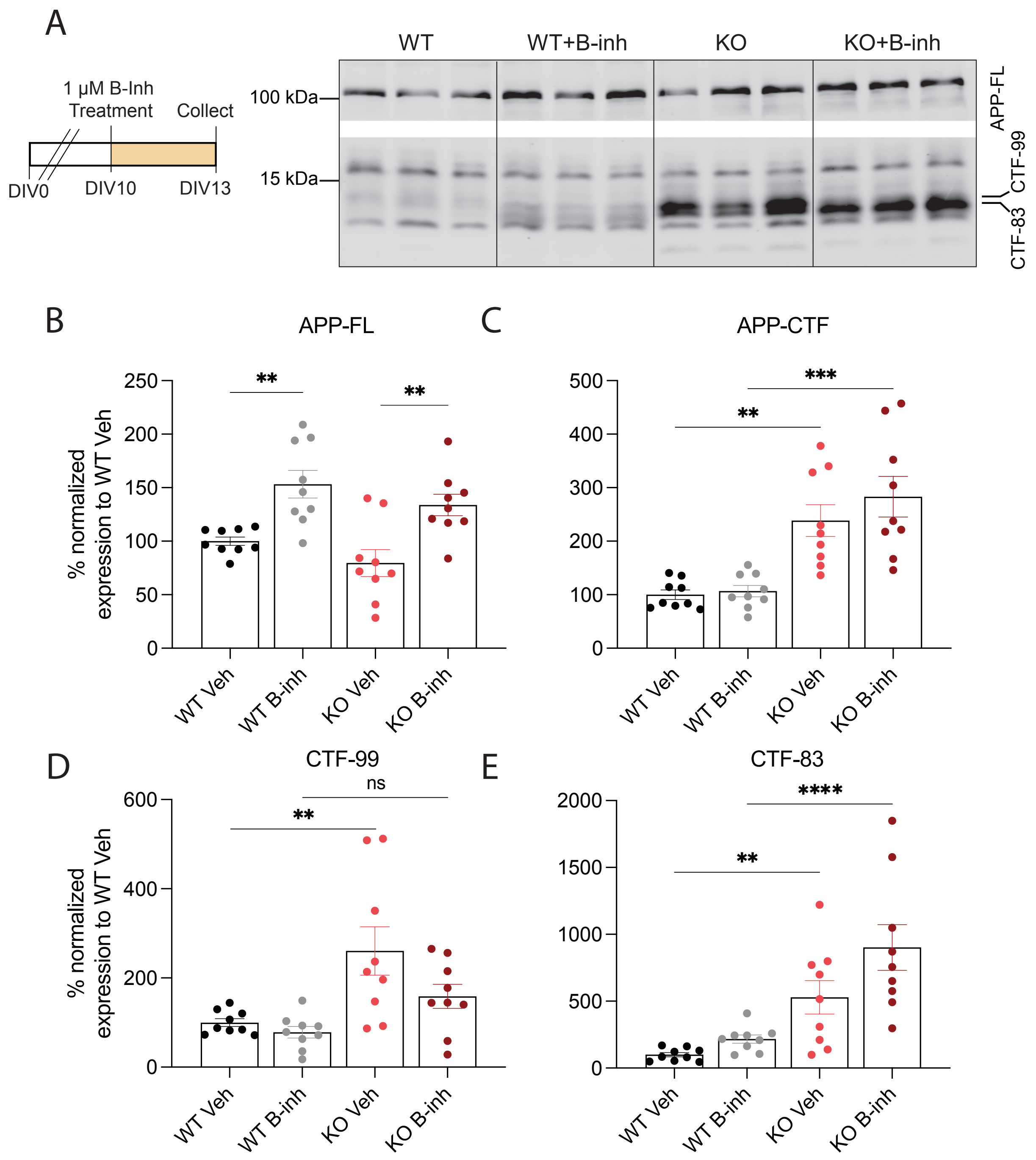

*Supplementary Figure 2*

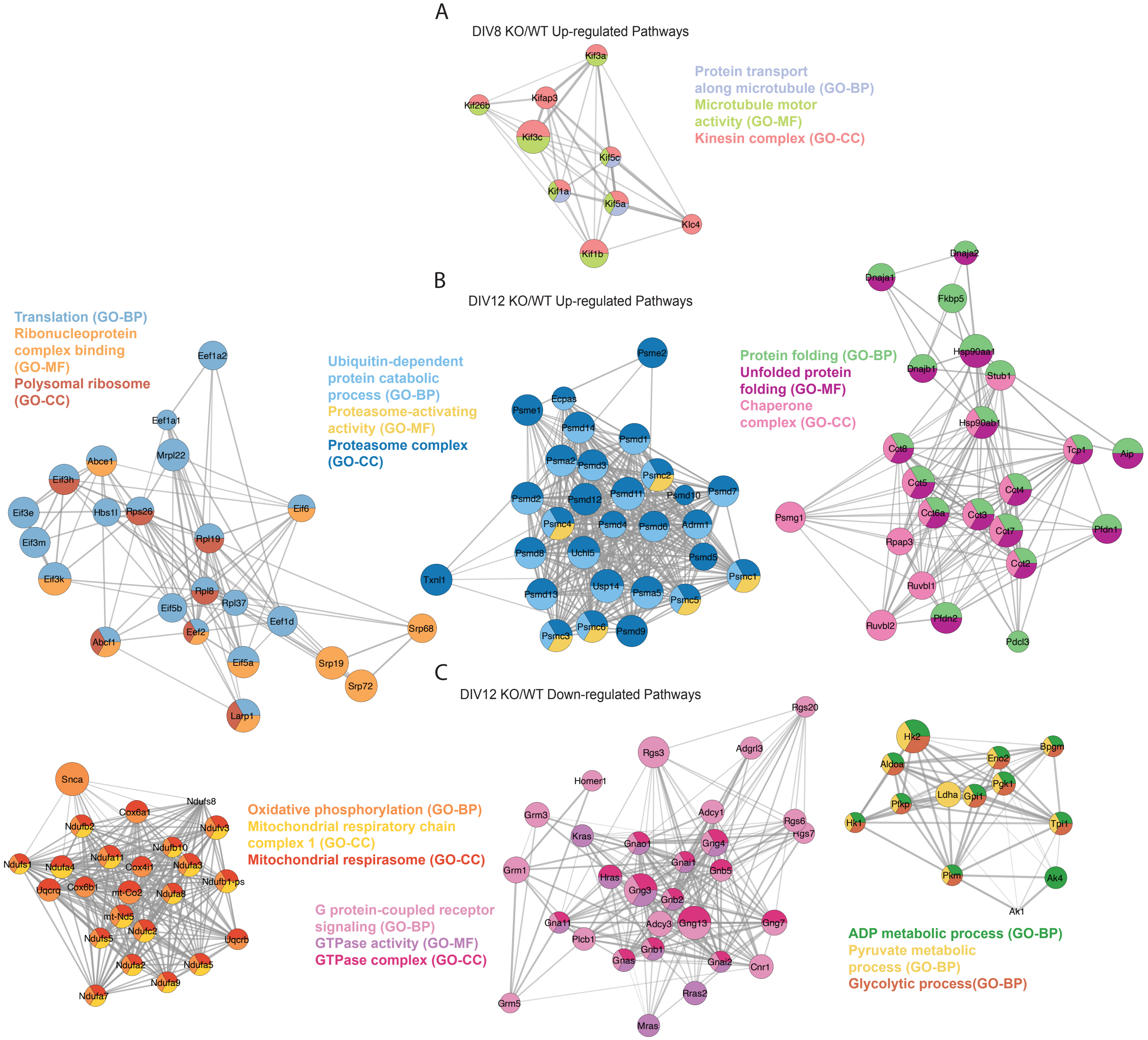
